# Studies on the co-metabolism of glucose and glycerol in the fungus *Umbelopsis isabellina*

**DOI:** 10.1101/2022.12.14.520399

**Authors:** Panagiotis Dritsas, George Aggelis

## Abstract

Over the past few years it is observed an increased interest for oleaginous microorganisms in the perspective to produce microbial oils of great commercial interest through the consumption of low/zero cost substrates. In this paper, the physiology of the fungus *Umbelopsis isabellina* growing on blends of glycerol and glucose was investigated. In all experiments the fungus completely consumed glucose and produced satisfactory quantities of biomass containing reserve lipids in high percentages. However, glycerol concentration in the growth medium was negatively correlated to glucose assimilation rate, mainly during the balanced-growth phase. Nevertheless, at high initial concentrations, glycerol was partially consumed and seemed to contribute positively to the suppression of lipid degradation. Following the discovery of this complex regulatory mechanism regarding glucose and glycerol co-assimilation, the activity of three key-enzymes namely aldolase, glycerol kinase and glycerol dehydrogenase, which are implicated in glycerol and glucose assimilation, was investigated. The experiments revealed a clear preference of the fungus for glucose over glycerol. On the other hand, storage polysaccharides are degraded instead of storage lipid at the late oleaginous phase for maintenance purpose. These new biochemical features will enable the design of appropriate growth media for the co-fermentation of these two substrates by *U. isabellina* with the aim to maximize lipid accumulation.

## 1. Introduction

*Umbelopsis isabellina* is a cosmopolitan heterotrophic saprophytic filamentous fungus which is considered a significant candidate for a wide range of industrial applications, i.e., biodiesel, pharmaceuticals and chemicals production (Crowther, Boddy and Hefin Jones 2012; Papanikolaou and Aggelis 2019). *U. isabellina* carries the capacity of growing over a wide range of environmental conditions, such as temperature, pH and oxygen concentrations (Ferreira *et al*. 2013; Wang *et al*. 2018), more importantly, though, it has the ability to metabolize various industrial substrates of low or even zero cost and synthesize metabolites of high-added value in significant quantities (Fakas *et al*. 2009; Chatzifragkou *et al*. 2010; Ruan *et al*. 2012; Demir *et al*. 2013; Zeng *et al*. 2013). For instance, *U. isabellina* can accumulate unusually high lipid amounts, i.e., up to 80% w/w in dry biomass, under specific growth conditions (Subramaniam *et al*. 2010). Its lipids contain γ-linolenic acid, a polyunsaturated fatty acid (PUFA) of importance for dietary and pharmaceutical purposes (Papanikolaou, Komaitis and Aggelis 2004; Bellou *et al*. 2012; Gao *et al*. 2013) while its more saturated fatty acids can be channeled to the biodiesel production industry (Meeuwse *et al*. 2012). Last but not least, its ability to degrade and/or reduce the toxicity of environmental pollutants (metals, phenolic compounds, etc.) has been reported (Janicki, Długoński and Krupiński 2018).

Despite *U. isabellina* was first described back in 1902 (Meyer and Gams 2003), its biology and ecology are still not well understood. Here we are interested in the ability of *U. isabellina* to co-assimilate glucose and glycerol, a feature of potential industrial interest. Most heterotrophic microorganisms are able to utilize a wide range of sugars, but usually glucose is the most preferable one. Notably, the presence of glucose in the growth medium suppresses the biosynthesis of enzymes involved in both the transport and the catabolic process of other sugars, phenomenon known as “glucose effect”. Assimilation of other sugars is usually activated only after glucose depletion in the growth medium (Brauer *et al*. 2005; Martínez-Gómez *et al*. 2012; Bommareddy *et al*. 2015). On the other hand, some oleaginous microorganisms can grow sufficiently on glycerol as the sole carbon source and accumulate PUFAs in significant quantities (Chatzifragkou *et al*. 2011; Siramon, Punsuvon and Riengsilchai 2016; Dourou *et al*. 2017; Mironov *et al*. 2018). Glycerol is a byproduct of biodiesel, soap and alcoholic beverage plants, which is considered an important low acquisition cost substrate for industrial microbiology (Nicol, Marchand and Lubitz 2012; Cai *et al*. 2019; Chmielarz *et al*. 2021).

In this study, *U. isabellina* was cultivated on growth media containing blends of glycerol and glucose as carbon sources at different concentrations. The ability of the fungus to assimilate glycerol in the presence of glucose during the various growth phases, such as the balanced growth phase and the lipid accumulation phase, the activity of key-enzymes during the catabolism of these two substrates, and its ability to accumulate intracellular lipids under these conditions were studied. The elucidation of the ways glucose affects the catabolism of glycerol and *vice versa* will enable the design of appropriate growth media for the co-fermentation of these two substrates, with the perspective maximizing their assimilation by *U. isabellina* towards maximum lipid production.

## 2. Materials and methods

### 2.1 Biological material and culture conditions

The fungus *Umbelopsis isabellina* ATHUM 2935 (culture collection of National and Kapodistrian University of Athens, Greece) was kept for maintenance purposes on potato dextrose agar (Biolab Zrt, Budapest, Hungary) at 7 ± 1°C and master cultures were regularly sub-cultured.

Growth media containing as carbon and energy sources glucose (AppliChem, Darmstadt, Germany) and/or glycerol (of purity 99.8%, Fluka, Steinheim, Germany), in various concentrations (see Table 1), were supplemented with minerals (in g L^-1^): KH_2_PO_4_ (AppliChem), 12.0; Na_2_HPO_4_ (AppliChem), 12.0; CaCl_2_.2H_2_O (Carlo Erba, Rodano, Italy), 0.1; CuSO_4_.5H_2_O (BDH, Poole, England), 0.0001; Co(NO_3_).6H_2_O (Merck, Darmstadt, Germany), 0.0001; MnSO_4_.5H_2_O (Fluka), 0.0001; ZnSO_4_.7H_2_O (Merck), 0.001 and FeCl_3_.6H_2_O (BDH), 0.08. The media were nitrogen-limited with yeast extract (Conda, Madrid, Spain) at 3.0 g L^-1^ being the sole nitrogen source, while it was also deployed as source of magnesium and ferrum, since the aforementioned composition has been proved to induce lipid accumulation (Bellou et al. 2016). Medium pH after sterilization at 121°C for 20 min was 6.5 ± 0.5 and remained practically stable during cultivation, due to the high buffer capacity of the media containing Na_2_HPO_4_ and KH_2_PO_4_. Glucose, Na_2_HPO_4_ and CaCl_2_ were sterilized separately to avoid medium alteration (Makri, Fakas and Aggelis 2010) and added into the flasks aseptically.

**Table 1.**
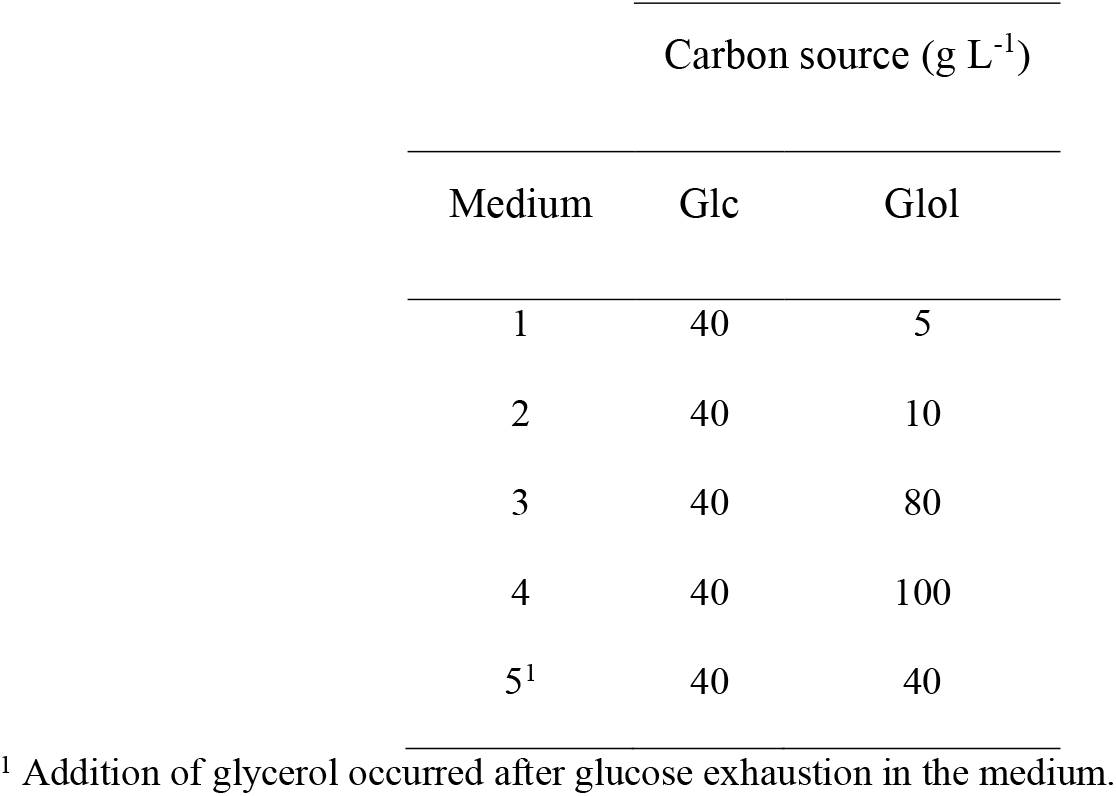
Concentrations of the carbon sources (i.e., glucose - Glc and glycerol - Glol) in the different culture media used in this study.

| Medium | Carbon source (g L<sup>-1</sup>) |  |
| --- | --- | --- |
|  | Glc | Glol |
| 1 | 40 | 5 |
| 2 | 40 | 10 |
| 3 | 40 | 80 |
| 4 | 40 | 100 |
| 5<sup>1</sup> | 40 | 40 |

<sup>1</sup> Addition of glycerol occurred after glucose exhaustion in the medium.

Submerged cultures were performed in 250 mL Erlenmeyer flasks containing 50 mL of growth medium. Flasks were inoculated with 1 mL of spore suspension containing 10^7^ fungal spores. Incubation was held in an orbital shaker (ZHICHENG ZHWY 211C, Shanghai, China), equipped with a ventilation system, at 28 ± 1°C and an agitation rate of 180 rpm.

### 2.2 Analytical methods

#### Cell mass determination

Flasks were removed from the incubator on a daily basis and fungal mycelia were harvested by filtration under vacuum through Whatman No. 1 paper. The collected cell mass was washed three times with distilled water and then dried at 80°C until constant weight. The final step was the gravimetrical determination of the total cell mass (x, g L^-1^). Lipid-free cell mass (x_f_) was calculated by subtracting cellular lipids (determined as described in the next subsection) from total cell mass.

#### Lipid extraction

Total mycelial lipids were extracted in chloroform: methanol 2:1 (v/v) (Sigma-Aldrich) according to the Folch, Lees and Sloane Stanley (1957) method and the extracts were filtrated through Whatman No. 1 paper. Following solvent evaporation under vacuum, using a Rotavapor R-20 device (BUCHI, Flawil, Switzerland), the total cellular lipid quantities were gravimetrically determined and expressed as the percentage of lipids in the dry cell mass (L/x%).

#### Fatty acid composition of cellular lipids

Total lipids were converted into their fatty acid methyl-esters (FAMEs) in a two-step reaction, in accordance with the AFNOR (1984) method, and analyzed with Gas Chromatography (GC) towards the investigation of fatty acid composition. A GC device (Agilent 7890A device, Agilent Technologies, Shanghai, China), equipped with a flame ionization detector at 280°C, using a HP-88 (J&W Scientific) column (60 m × 0.32 mm) was used for the analyses. Carrier gas was helium at a flow rate 1 mL/min and the analysis was run at 200°C. Peaks of FAMEs were identified through comparison to authentic standards.

#### Glucose and glycerol determination

Glucose and glycerol concentrations were determined in filtered aliquots of the culture supernatant by High Performance Liquid Chromatographer (HPLC). Specifically, the HPLC apparatus (Ultimate 3000 Dionex, Germering, Germany) was equipped with an Aminex HPX-87H column (300 × 7.8 mm) as well as UV-Vis and RI detectors. HPLC analyses were conducted under the following conditions; mobile phase: H_2_SO_4_ (Fluka) 0.004 N, flow rate: 0.9 mL min^-1^ and column temperature: T = 55°C.

#### Microscopy

Mycelial morphology was observed using a Carl Zeiss (GmbH, Gottingen, Germany) optical microscope, equipped with a digital video camera (Exwave HAD, Sony, Tokyo, Japan), on a daily basis.

The intracellular lipid bodies were stained with Nile red (Sigma-Aldrich) fluorescence dye following Greenspan, Mayer and Fowler (1985) method, and visualized under an Axiostar 40 (Zeiss, Cambridge, UK) fluorescence microscope equipped with an excitation filter of 470/40 nm and a ProgRes camera (Jenoptik CF cool, Jena, Germany).

#### Determination of enzymatic activities

##### Cell extract preparation

Harvested fungal mycelia of known culture volume were washed twice with a 50 mM Na_2_PO_4_/KH_2_PO_4_ pH 7.5 buffer and then resuspended in a 30 mM Na_2_PO_4_/KH_2_PO_4_ pH 7.5 buffer, which contained 1 mM DTT (Sigma -Aldrich), 1 mM benzamidine (Fluka) and 250 mM sucrose (Merck), at a ratio 1 mL buffer: 0.5 g wet cell mass. Consequently, the cells were ruptured by three 2 min sonic bursts of 90 W and one 1 min sonic burst of 90W at 0–4°C using a sonicator device (Sonics Vibra cell CV188, Newtown, CT, USA). Afterwards, the disrupted cells were centrifuged at 24.000g for 50 min at 4°C and the supernatant was filtered through a Whatman 0.2 μm membrane in order to remove solidified lipids and other remaining cell debris.

##### Enzyme assays

Aldolase (ALDO, EC 4.1.2.13) was assayed according to the method described by Bergmeyer (1974), by measuring ΔA_340_ as the result of β-NADH oxidation. The activity of glycerol kinase (GK, EC 2.7.1.30) was measured at ΔA_366_ as the result of NAD^+^ reduction, following Bublitz and Kennedy (1954) method. Finally, glycerol dehydrogenase (GDH, EC 1.1.1.6) activity was measured in accordance with Burton (1955) method, by measuring ΔA_340_ which is dependent on the β-NAD reduction. The enzymatic activities were determined in a Shimadzu 1800 UV-Vis spectrophotometer at 25°C.

One unit (U) of enzyme was defined as the amount of enzyme catalyzing the formation of 1 μmol of each enzymatic reaction product/min under the conditions of the aforementioned methods. Soluble protein was determined according to Papanikolaou et al. (2004), and specific activity was expressed as units per gram of soluble protein.

#### Statistical analysis

The experimental data from *U. isabellina* cultures and growth kinetics were treated using OriginPro 2021 9.8.0.200 ®, 1991–2020. Two independent cultures were performed for enzyme activities (units/g of fat-free biomass) determination in cell-free extract sample of *U. isabellina* and are presented as mean values from at least two replications.

## 3. Results

### 3.1 Kinetics of growth and lipid accumulation

Kinetics of growth (in terms of dry fat-free biomass - x_f_, g L^-1^), lipid accumulation (L/x%) and glucose (Glc, g L^-1^) consumption, were investigated regarding the different growth media. It was found that *U. isabellina* cultivated on blends of glucose 40 g L^-1^ and glycerol at low concentrations (i.e. 5 g L^-1^ and 10 g L^-1^) totally assimilated glucose after approximately 120 h of cultivation (Fig. 1a), while consuming very low glycerol amounts (data not shown). The addition of glycerol at high concentrations (i.e. 80 g L^-1^ and 100 g L^-1^) in the growth medium, caused a delay in glucose consumption, especially in the first hours of incubation (Fig. 1a). As for glycerol assimilation, it amounted to 6.3 g L^-1^ and 12.8 g L^-1^, for glycerol initial concentration 80 and 100 g L^-1^, respectively (data not shown).

**Figure 1.**
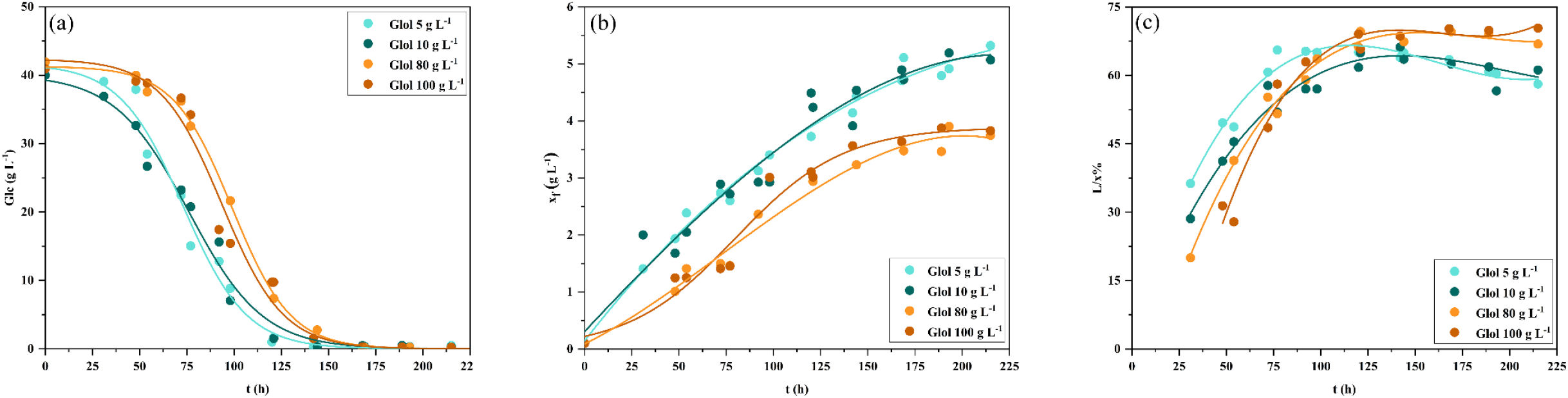
Kinetics of (a) glucose consumption (Glc, g L^-1^), (b) dry fat-free biomass (x_f_, g L^-1^) production and (c) lipid accumulation (L/x%, w/w) in the mycelia of *U. isabellina* growing in the presence of glycerol (Glol) as additional carbon source at various concentrations (media 1-4 shown in Table 1).

Fat-free biomass production was favored in low glycerol content media and was around 5.3 g L^-1^ (Fig. 1b). In these media the accumulation of intracellular lipids was maximized at 120 h, reaching L/x around 65.1%, followed by partial lipid degradation to L/x around 58.1% at 215 h (Fig. 1c). On the other hand, when glycerol was added at higher initial concentrations in the growth medium (i.e. 80 g L^-1^ and 100 g L^-1^), the maximum fat-free biomass production significantly reduced to around x_f_ = 4 g L^-1^. On the contrary, lipid accumulation was slightly higher comparing to the experiments with low glycerol concentrations, reaching the values of L/x = 70%, though lipid degradation was not observed in both cases. The lipid droplets morphology and the chlamydospores of *U. isabellina* after Nile Red staining under fluorescence microscopy are depicted in Fig. 2.

**Figure 2.**
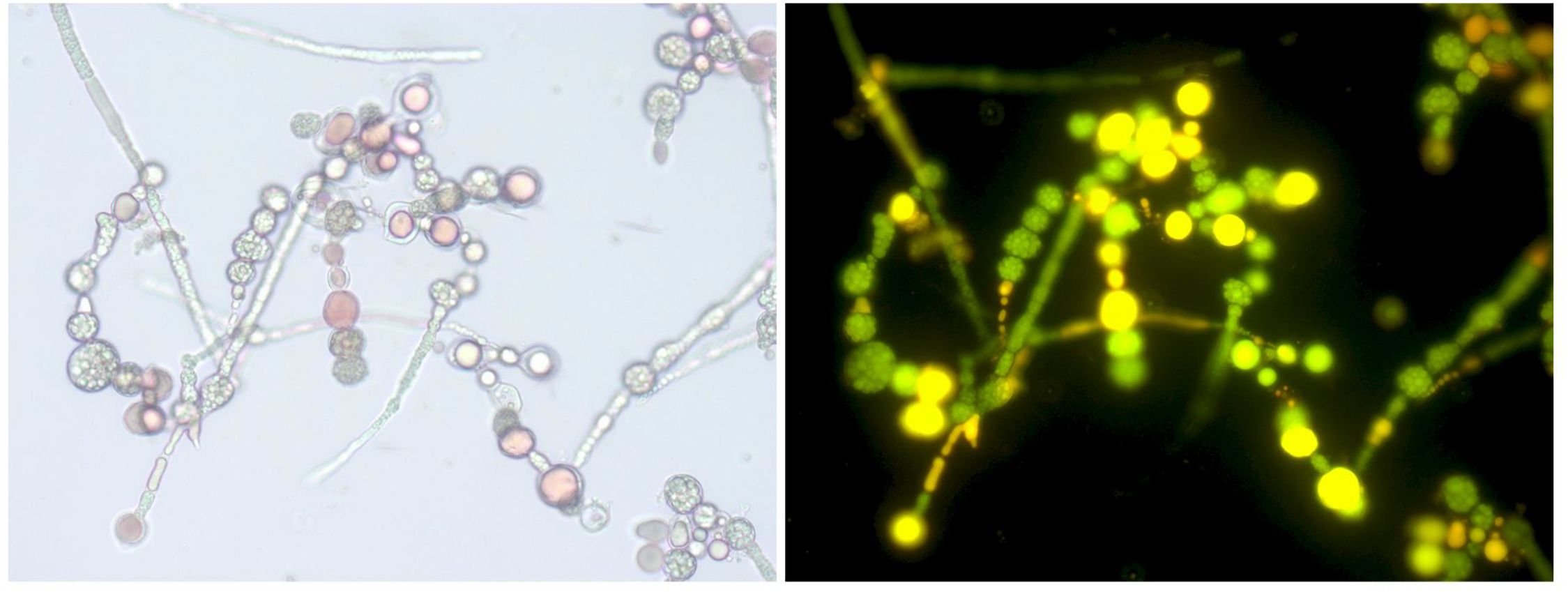
Lipid droplets morphology in the mycelia and the chlamydospores of *U. isabellina* under optical (on the left) and fluorescence (on the right) microscopy after Nile Red staining at 138 h of cultivation in Medium 2 of Table 1.

When glycerol was added in the growth medium (at 40 g L^-1^), upon complete consumption of glucose, a consumption of glycerol occurred, in parallel with repression, partially at least, of lipid degradation (Fig. 3).

**Figure 3.**
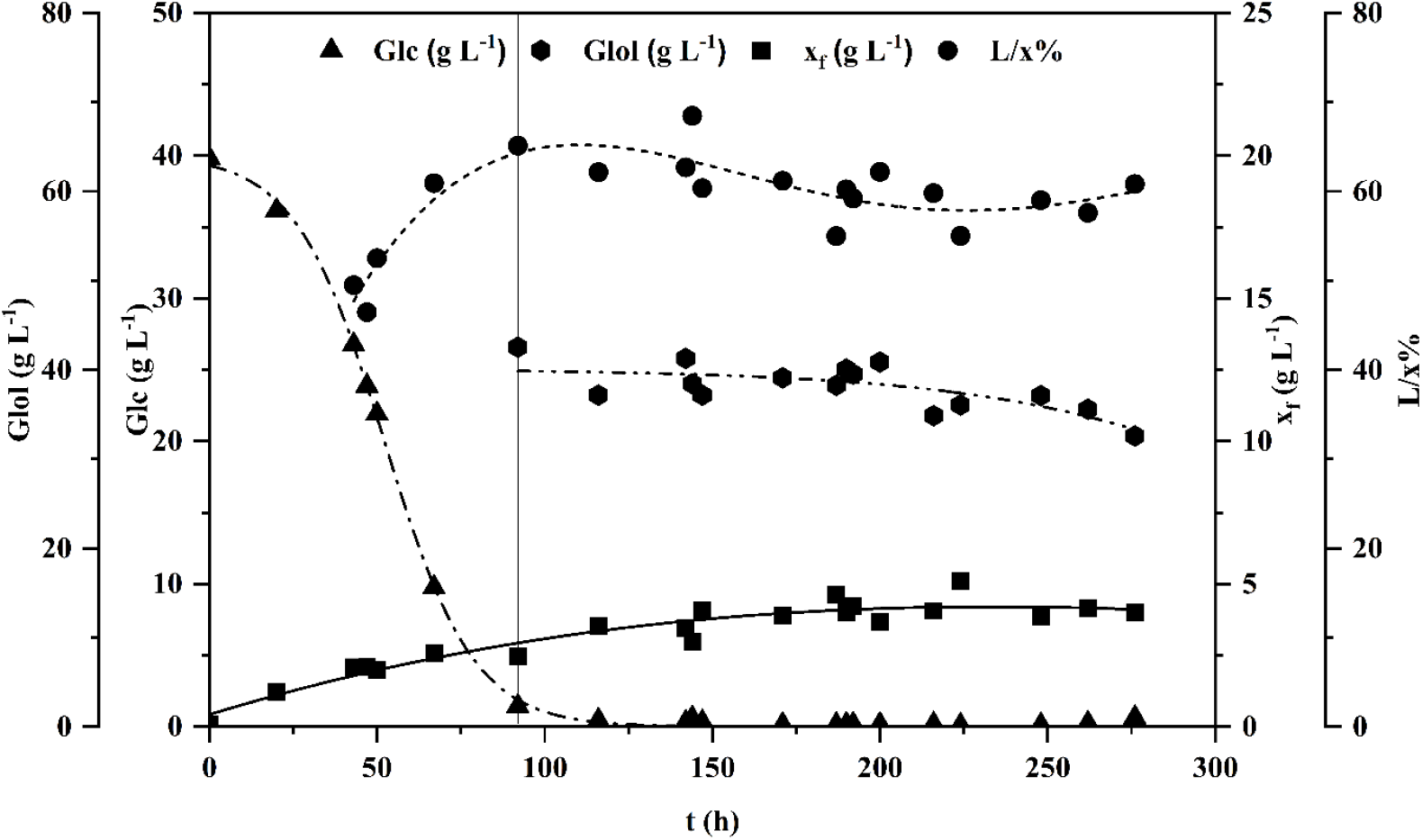
Growth kinetics of glucose (Glc, g L^-1^) and glycerol (Glol, g L^-1^) consumption, total dry biomass (x, g L^-1^) production and lipid accumulation (L/x%, w/w) in the mycelia of *U. isabellina*. The vertical line indicates the time of glycerol (Glol) addition at 40.0 g L^-1^ in the growth medium (t = 92 h). Culture conditions: mineral medium (see Material and Methods section) with Glc 40.0 g L^-1^ as carbon sources and yeast extract at 3.0 g L^-1^ as source of nitrogen, magnesium and growth factors; pH = 6.5 ± 0.5; temperature 28.0 ± 1.0°C; agitation rate 180 rpm.

### 3.2 Fatty acid composition

The fatty acid composition of total lipid of *U. isabellina* at the time of maximum lipid accumulation in the different culture media are gathered in Table 2. Although some small differences can be observed, no specific trend in fatty acid composition could be established. The main cellular fatty acids produced were oleic acid (^Δ9^C18:1) (percentage range of 54.0-63.5%, w/w), palmitic acid (C16:0) (17.9-24.3%, w/w) and linoleic acid (^Δ9,12^C18:2) (10.9-12.9%, w/w). The fatty acids stearic (C18:0), palmitoleic acid (^Δ9^C16:1) and gamma-linolenic acid (^Δ6,9,12^C18:3) were found in lower percentages.

**Table 2.**
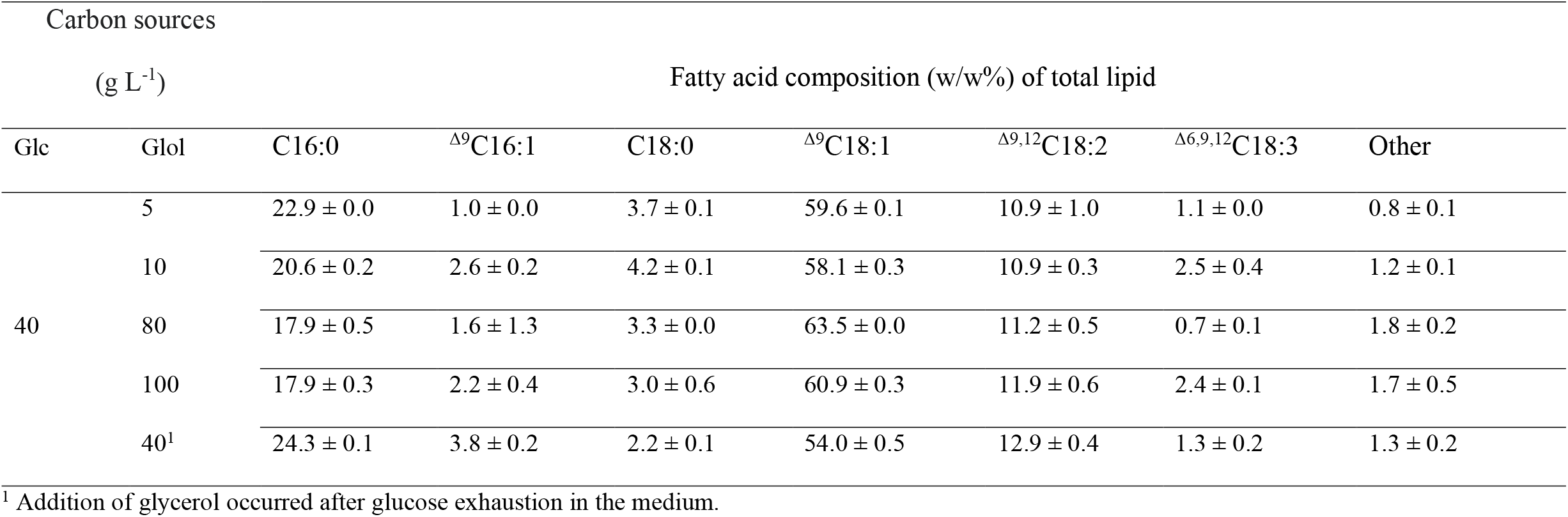
Fatty acid composition of total lipid of *U. isabellina* at the time of maximum lipid accumulation in the different culture media. Culture conditions: Mineral medium (see Material and Methods section) with yeast extract at 3.0 g L^-1^ as source of nitrogen, magnesium and growth factors; pH = 6.5 ± 0.5; temperature 28.0 ± 1.0°C; agitation rate 180 rpm.

### 3.3 Enzyme activity

The activities of Glycerol Kinase (GK), Glycerol Dehydrogenase (GDH) and Aldolase (ALDO) of cell-free extract of *U. isabellina* cultivated under different concentrations and combinations of glucose and glycerol, at selected physiological phases were determined (Table 3). Specifically, when the fungus was cultivated on blends of glucose 40 g L^-1^ and glycerol 5 g L^-1^ or 100 g L^-1^, enzyme activities were determined in the balanced growth phase (at 48 h) and in the late oleaginous phase (at 180 h), while in the case that addition of glycerol in the medium took place after glucose exhaustion, the selected checkpoints were 120 h (completion of the oleaginous phase) and 260 h (potential phase of degradation). In the cases of glucose and glycerol blends, GK activity was in the balanced growth phase higher compared to the late oleaginous phase, while in the experiment that glycerol addition to the medium followed glucose exhaustion, was low during both phases. GDH activity showed similar trend with GK at the selected time checkpoints as shown in Table 3. When growth occurred in the blends’ medium, ALDO activity was very high in the balanced growth phase, but dramatically drops in the late oleaginous phase. However, when glycerol was added later on the medium, ALDO activity was significantly increased (from 3.9 ± 1.0 to 20.5 ± 0.1 U/g(x_f_)_260h_).

**Table 3.** Enzyme activity of Glycerol Kinase (GK), Glycerol Dehydrogenase (GDH) and Aldolase (ALDO) of cell-free extract (U/g_(_x_f)_) of *U. isabellina* cultivated under different concentrations and combinations of glucose (Glc) and glycerol (Glol), at selected culture times. The produced dry biomass, fat-free biomass and total lipids content of the aforementioned times are presented.

| Carbon source | | Growth phase | x<br>(g L<sup>-1</sup>) | L<br>(g L<sup>-1</sup>) | L/x<br>(%) | x<sub>f</sub><br>(g L<sup>-1</sup>) | GK<br>$U/g(x_f)$ | GDH<br>$U/g(x_f)$ | ALDO<br>$U/g(x_f)$ |
| --- | --- | --- | --- | --- | --- | --- | --- | --- | --- |
| Glc | GloI<br>(g L<sup>-1</sup>) |  |  |  |  |  |  |  |  |
| 40 | 5 | Early oleaginous (t = 48 h) | 4.0 ± 0.2 | 2.0 ± 0.1 | 50.8 ± 1.2 | 2.0 ± 0.0 | 22.0 ± 0.0 | 59.0 ± 0.9 | 119.8 ± 3.8 |
|  |  | Late oleaginous (t = 180 h) | 12.1 ± 0.2 | 7.8 ± 0.4 | 64.7 ± 3.9 | 4.3 ± 0.5 | 3.6 ± 0.9 | 5.9 ± 0.4 | 12.8 ± 0.6 |
|  | 100 | Early oleaginous (t = 48 h) | 2.5 ± 0.8 | 0.8 ± 0.3 | 29.7 ± 1.8 | 1.7 ± 0.5 | 67.3 ± 1.8 | 31.4 ± 1.3 | 209.2 ± 2.7 |
|  |  | Late oleaginous (t = 180 h) | 12.5 ± 0.3 | 8.8 ± 0.2 | 70.0 ± 0.2 | 3.8 ± 0.1 | 19.4 ± 0.4 | 5.3 ± 1.3 | 29.9 ± 2.1 |
|  | 40<sup>1</sup> | Late oleaginous (t = 120 h) | 9.3 ± 0.1 | 6.7 ± 0.5 | 71.9 ± 5.0 | 2.6 ± 0.4 | 2.9 ± 0.4 | 3.5 ± 0.0 | 3.9 ± 1.0 |
|  |  | Potential lipid turnover (t = 260 h) | 10.0 ± 0.2 | 6.2 ± 0.6 | 62.2 ± 4.6 | 3.8 ± 0.4 | 3.6 ± 0.3 | 4.9 ± 0.4 | 20.5 ± 0.1 |

<sup>1</sup> Addition of glycerol occurred after glucose exhaustion in the medium

## 4. Discussion

Purpose of this study was to investigate the ability of *U. isabellina* to co-assimilate glucose and glycerol during the different phases of its growth cycle. Glucose is the most commonly used carbon substrate in Microbiology, while glycerol, as an important by-product of various industrial processes, is a potential carbon source for chemical and biotechnological processes. In fact, the conversion of glycerol into valuable chemicals through microbial processes is a modern and quite efficient biotechnological approach of its utilization which circumvents the disadvantages of chemical conversions.

Regarding microbial cultures on glucose and glycerol used as co-substrates, two different trends of carbon uptake were observed so far. The first one is the classic diauxic growth, observed for instance, when *Escherichia coli* or *Rhodosporidium toruloides* were cultivated on blends of glucose and glycerol (Martínez-Gómez *et al*. 2012; Bommareddy *et al*. 2015). In both cases, glucose was consumed primarily but upon its depletion from the media, the assimilation mechanisms of glycerol were activated. The second one is the simultaneous assimilation of these two substrates, observed in the yeasts *Yarrowia lipolytica* and *Rhodotorula glutinis* (Easterling et al. 2009; Workman, Holt and Thykaer 2013). In contrast to the above, interesting interactions in glucose and glycerol uptake were observed in all experiments performed in the current investigation. Specifically, glucose exhaustion in high glycerol concentration media (i.e. 80 g L^-1^ and 100 g L^-1^) occurred approximately 30 h later compared to media containing glycerol in low concentrations, suggesting inhibition of glucose uptake by glycerol. The inhibitory role of glycerol to glucose uptake, especially during balanced-growth phase, could justify the partial suppression of x_f_ production under high concentrations of glycerol in the medium. In contrast, lipid accumulation appeared to be unaffected by the presence of glycerol in the medium. As for glycerol assimilation, when added at low concentrations to the medium it was not taken up, while partial glycerol uptake was observed when added at high concentrations, probably via simple diffusion. Limited glycerol assimilation occurred as well when it was added after glucose depletion in the medium. The aforementioned show that *U. isabellina*, exhibits a clear preference for glucose, a feature that appears to be common among the Mucorales (Papanikolaou, Komaitis and Aggelis 2004; Papanikolaou et al. 2007; Chatzifragkou et al. 2010; Meussen et al. 2012; Kowalczyk et al. 2018). Furthermore, it is established that Mucorales show poor assimilation ability for glycerol (Papanikolaou et al. 2010, 2017; Chatzifragkou et al. 2011) with the fungus *Mortierella ramanniana* MUCL 9235 being the only recorded exception (Bellou et al. 2012).

Glycerol used as co-substrate in high concentrations, in spite of its low consumption, represses reserve lipid turnover which usually occurs after glucose exhaustion in the growth medium. This finding was confirmed by an additional experiment in which glycerol, added to the medium after glucose depletion, suppressed the consumption of the lipid reserves. Furthermore, it must be noted that in high glycerol content media, lipid accumulation levels were slightly enhanced. Likewise, when *U. isabellina* ATCC 42613 was cultured on pretreated bagasse enzyme hydrolysate (containing glucose and glycerol), a positive correlation between glycerol concentration and oil content was recorded (Cai et al. 2019). Specifically, the increase of glycerol content in the medium from 0% to 2.0% led to a slight increase in oil content from 49.6% to 52.3%. The authors attributed this trend to the high C/N ratios which promote lipid accumulation. Thus, glycerol seems to be a suitable carbon source for suppressing turnover of lipid reserve. The fatty acid profile of lipids produced in the presence of glycerol was similar to that of lipids produced by *U. isabellina* grown on glucose and similar sugars (Chatzifragkou et al. 2010; Vamvakaki et al. 2010; Papanikolaou et al. 2017; Cai et al. 2019).

The aforementioned data that were abstracted from the kinetics of Figs. 1 and 3 revealed an intriguing regulatory mechanism regarding glucose catabolism in the presence of glycerol and *vice versa* which affects biomass production and lipid accumulation. In the aftermath of this emerging complex regulatory mechanism, the activity of certain key-enzymes implicated in glycerol (GK and GDH) and glucose (ALDO) assimilation was studied. In the early oleaginous phase ALDO noted significant levels of activity, confirming the clear preference of *U. isabellina* for glucose over glycerol. Besides, Clapés *et al*. (2010), experimenting with *E. coli*, proved that ALDO functions as an allosteric inhibitor of GK in gluconeogenesis, inhibiting glycerol catabolism. In contrast, when glycerol was added in the growth medium, upon complete consumption of glucose, ALDO activity presented increase at the potential lipid degradation phase. When Dourou *et al*. (2017) cultured *U. isabellina* ATHUM 2935 (i.e. the strain of the present study) on a growth medium with glucose as the sole carbon source, observed that after glucose depletion, the accumulated polysaccharides were utilized as an intracellular energy source for maintenance purposes or converted to lipids. Therefore, we can attribute the high values of ALDO activity in degradation/turnover of storage polysaccharides, which may contribute, additionally to glycerol consumption, to the repression of lipid turnover.

The activity of GK was more intense during the early oleaginous phase, especially when the fungus grew on the blend with initial concentrations of glucose 40 g L^-1^ and glycerol 100 g L^-1^. Several reports establish the critical importance of GK for glycerol assimilation, despite the existence of alternative pathways for glycerol catabolism. For example, mutant strains of *E. coli*, which were unable to synthesize GK, exhibited growth failure on glycerol, despite the presence of the genes encoding glycerol dehydrogenase and dihydroxyacetone kinase (Lin 1976; Sprenger 2017). Moreover, when Hao *et al*. (2015) overexpressed the genes coding GK to improve the glycerol assimilation capacity in the fungus *Mortierella alpina*, there was an increase (>35%) in the fatty acids produced compared to the control experiment. Our results indicate a preferred kinase pathway for glycerol assimilation for *U. isabellina* as well.

Finally, the trend of GDH activity in the experiments was similar to that of GK. Izawa *et al*. (2004) observed that the removal of GDH coding genes led to increased intracellular glycerol accumulation in *Saccharomyces cerevisiae*. Consequently, GDH plays a crucial role in the synthesis and final production of glycerol intracellularly, contributing to the regulation of osmotic stress. *S. cerevisiae* appeared to synthesize glycerol intracellularly in response to nitrogen starvation conditions as well. The addition of nitrogen sources, like amino acids, peptone or yeast extract affected positively the capacity of *S. cerevisiae* to grow on glycerol (Merico *et al*. 2011; Swinnen *et al*. 2013). However, there are also microorganisms where the presence and activity of GDH in glycerol assimilation is considered to be of high importance, such as in the yeast *Schizosaccharomyces pombe* (Matsuzawa *et al*. 2010).

The conversion of glycerol into the microbial cell is regulated by several factors. For a long time, glycerol was regarded as a molecule which permeates a plasma membrane solely by passive diffusion, driven by a concentration gradient (Gancedo, Gancedo and Sols 1968; Romano 1986). However, recent data overthrew the above hypothesis. For instance, the product of an already identified gene (i.e., STL1 gene) in the yeasts *S. cerevisiae* and *Candida albicans* (Ferreira *et al*. 2005; Kayingo *et al*. 2009) which was recently detected by da Cunha *et al*. (2019) in the fungus *Wickerhamomyces anomalus*, functions as a glycerol/H^+^ symporter. Interestingly, the expression of this gene was inhibited in the presence of glucose, whereas the presence of glycerol, in relatively high concentrations, activated it. Additionally, when Ferreira *et al*. (2005) removed this gene from *S. cerevisiae*, the active transport of glycerol into the cells was not possible. These findings support the assumption that the activity of GK and GDH in our experiments may be due to the catabolism of intracellular glycerol produced during cell metabolism. It must be noted, though, that other genes, such as GUP1 and GUP2, which encode membrane proteins with multiple domains, probably function indirectly on glycerol uptake (Neves, Lages and Lucas 2004; Oliveira and Lucas 2004; Bosson, Jaquenoud and Conzelmann 2006).

## Conclusions

The obtained results revealed the existence of a complex regulation mechanism of glycerol and glucose co-assimilation in the fungus *U. isabellina*. When glycerol was present in the medium, glucose assimilation was delayed mainly during balanced-growth phase, and therefore partially suppressed x_f_ production. However, glycerol seemed to contribute suppress lipid degradation, especially when used in high concentrations. Thus, it can be used as a carbon source during the late oleaginous phase after glucose exhaustion, to suppress lipid turnover without altering fatty acid composition.

Meanwhile, the definite preference of *U. isabellina* to glucose was confirmed by the high values obtained regarding ALDO activity during early oleaginous phase and upon complete consumption of glucose. Surprisingly, ALDO activity was increased passing from the late oleaginous to the potential lipid degradation phase when the fungus was grown solely on glucose, even if glycerol was added after glucose depletion. This increase can probably be attributed to the degradation/recycling (turnover) of storage polysaccharides. The aforementioned is also supported by the relatively low activity of GK, which is usually directly connected to glycerol consumption. GDH function may be due to the response of cells to osmotic stress induced by extracellular glycerol or the catabolism of the intracellularly produced glycerol as a result of the fungus metabolism. However, the possibility of indirect factors that can influence the rate of glucose entry into the cells because of presence of glycerol cannot be excluded.

## Author agreement

Panagiotis Dritsas and George Aggelis have all agreed to submission.

## Acknowledgments

The project was supported by the University of Patras. The authors, therefore, acknowledge with thanks the University technical and financial support.

